# Adaptive radiation along ecological and morphological lines of least resistance in *Cyprinodon* pupfishes

**DOI:** 10.1101/2025.08.13.670168

**Authors:** Christopher H. Martin, Andrés Alvarez Zapata, Reyna Guadalupe Cetz Paredes, Frida Cortés, Sonia Gabriela Hernández, Matthew C. Kustra, Adan Fernando Mar-Silva, Fernando Mex, M. Fernanda Palominos, Charles Tralka, Maribel Badillo-Alemán, Juan J. Schmitter-Soto, Christopher M. Martinez, Jairo Arroyave, Carlos A. Gracida-Juárez

## Abstract

Adaptive radiation results in part from ecological opportunity in a new environment, but it is unclear how pre-existing constraints in the founding population may limit this process. Genetic lines of least resistance, and by proxy morphological variance, are known to limit adaptive divergence, but ecological variance is rarely investigated. Here we test whether ecological or morphological lines of least resistance in generalist populations may have constrained the directions of species divergence in two independent Caribbean adaptive radiations of *Cyprinodon* pupfishes. We find almost universal congruence between the major multivariate dimensions of intraspecific craniofacial and dietary variance within generalist populations and the major axes of interspecific divergence within each adaptive radiation. This is surprising given that we document unique trophic specialists within each radiation, including a bivalve-specialist, zooplanktivore, molluscivore/ostracod-specialist, and scale-eating specialist, while nearly all generalist populations were observed to feed rarely on these same resources. We conclude that pre-existing genetic constraints within each founding generalist population, resulting in dimensions of greater ecological and morphological variance, may partially constrain and predict the directions of species divergence and dietary specialization during adaptive radiation. We also provide a new framework for examining ecological lines of least resistance.

## Introduction

Adaptive radiation results from a rapid burst of adaptation and speciation. This process can result from the colonization of a new environment, evolution of a key innovation, or following a mass extinction, but a more quantitative framework for predicting adaptive radiation remains elusive (Martin & Richards, 2019; Stroud & Losos, 2020; Gillespie *et al*., 2020; De-Kayne *et al*., 2024) but see (Wagner, Harmon, & Seehausen, 2012, 2014). One highly influential hypothesis is that adaptive radiation should occur along genetic lines of least resistance: species divergence should be biased in the direction of the greatest multivariate additive genetic variance (the genetic variance-covariance matrix, *G*) within the ancestral founder population (Schluter, 1996, 2024). Empirical tests of this hypothesis often rely on measuring the principal components of the phenotypic variance-covariance matrix (*P*) as a proxy for G within one or more closely related outgroup populations, which serve as a proxy for the ancestral founding population. Despite strong evidence that the *G* matrix can evolve rapidly within a population over a few generations (Turelli, 1988; Roff, 2000; Steppan, Phillips, & Houle, 2002; Arnold *et al*., 2008), there is also substantial empirical evidence that radiations do tend to occur along the directions of greatest variance within *G* (or *P* as a proxy), even over million-year timescales (Martin, 2012; McGlothlin *et al*., 2018; Silva, Potts, & Harrison, 2020; Evans *et al*., 2021; Dunker, St John, & Martin, 2024; Schluter, 2024), but also see (Berner, Stutz, & Bolnick, 2010). Importantly, the topography of the adaptive landscape can also shape the *G* matrix, so these studies may indicate either genetic constraints on adaptive evolution or environmental selective pressures shaping the major dimensions of additive genetic variation during the process of radiation (Barton & Turelli, 1987; Eroukhmanoff, Eroukhmanoff, & Svensson, 2011).

The genetic lines of least resistance hypothesis can be extended to include lines of least resistance through axes of ecological diversity. Major dimensions of ecological variation within an ancestral population may be more easily co-opted for species divergence during adaptive radiation. This would result in major axes of intraspecific ecological variance significantly associated with major axes of species divergence within the radiation. Alternatively, the more traditional view is that ecological opportunities in a new environment may outweigh pre-existing major dimensions of variation in the founding population and bias species divergence in the direction of novel niche opportunities (Yoder *et al*., 2010; Gillespie *et al*., 2020). These possibilities are not mutually exclusive and may be difficult to separate, particularly if temporal shifts in environmental selective pressures also shape the outgroup populations used as a proxy for the ancestral population in parallel with the focal radiation. Nonetheless, parallel ecological divergence due to shared environmental selective pressures can be minimized by focusing on unique trophic specializations less affected by global environmental changes.

The hypothesis that ecological lines of least resistance bias species divergence during adaptive radiation has not been directly tested to our knowledge (but see (Deng *et al*., 2024). Traditionally, adaptive radiation is predicted to occur following an increase in ecological diversity within a population (Simpson, 1944; Dieckmann & Doebeli, 1999; Yoder *et al*., 2010) but without considering the initial major dimensions of multivariate niche diversity. There are many tests of trait utility (Schluter, 2000), the association between morphology and ecology within a radiation or within a single population, and substantial evidence of trait utility in natural populations (Bolnick & Lau, 2008; Bolnick & Paull, 2009; Kaeuffer *et al*., 2012a; Beausoleil *et al*., 2023). Similarly, inter-individual variation in niche breadth or individual niche specialization can also vary across populations (Bolnick, 2004; Svanbäck & Persson, 2004) and may promote radiation (Bolnick *et al*., 2007; Knudsen *et al*., 2010; Baguette *et al*., 2020). Nonetheless, it remains unclear whether the major dimensions of ecological diversity within the ancestral founding population can bias the direction of species divergence during adaptive radiation.

Here we tested if species divergence occurs along ecological and morphological lines of least resistance using two independent adaptive radiations of *Cyprinodon* pupfishes. Both pupfish radiations occur in sympatry within large inland lakes, each including endemic trophic specialists and a generalist omnivore (Martin & Wainwright, 2011). The Chichancanab Lake radiation in Mexico historically contained seven closely related species endemic to a 20-km long brackish lake (approximate salinity less than 3 ppt) in the center of the Yucatán peninsula (Humphries & Miller, 1981; Humphries, 1984; Strecker, 2002, 2005, 2006a). This included the generalist/detritivore Blackfin Pupfish *C. beltrani* and three trophic specialists: a predator that included adult fish in its diet, the Maya Pupfish *C. maya* (likely extinct), the crustacean and snail-eating benthivore Thicklip Pupfish *C. labiosus,* and the zooplanktivore Boxer Pupfish *C. simus* (Stevenson, 1992; Horstkotte & Strecker, 2005). Both *C. labiosus* and *C. simus* were collected for this study in 2022 and now appear to be extinct in the wild due to the recent invasion of the Mayan Cichlid *Mayaheros urophthalmus* (2022, 2024, 2025 pers. obs.) after the initial introduction of aquacultured Mozambique tilapia *Oreochromis mossambicus x niloticus* (Gracida-Juárez, Schmitter-Soto, & Genner, 2024) and colonization by native *Astyanax cf. bacalarensis* and *Poecilia velifera* in the early 1990’s (Schmitter-Soto & Caro, 1997; Schmitter-Soto, 1999; Fuselier, 2001; Strecker, 2006b; Gracida-Juárez *et al*., 2024).

The San Salvador Island (SSI) radiation in the Bahamas contains three closely related trophic specialist species endemic to the hypersaline lakes on this 21-km long island, including a specialized scale-eater *C. desquamator*, a molluscivore *C. brontotheroides* (Martin & Wainwright, 2013a), and an intermediate scale-eater *C*. sp. ‘wide-mouth’ (Richards & Martin, 2022; Palominos, Muhl, & Martin, 2024), which was not included in this study, in addition to the generalist/detritivore *C. variegatus* (Martin & Wainwright, 2013b; Hernandez *et al*., 2018). The SSI specialist species are most closely related to SSI *variegatus* and the radiation is nested within Caribbean *C. variegatus* populations, although there is clear evidence of secondary gene flow into the radiation from other regions of the Caribbean (Martin & Feinstein, 2014; Richards & Martin, 2017; McGirr & Martin, 2020, 2021; Richards *et al*., 2021; St John *et al*., 2024).

We used the most closely related generalist populations of *C*. *variegatus* and *C*. *beltrani* as a proxy for the ancestral founding population of each radiation, respectively (both species are primarily detritivores and omnivores consuming microcrustaceans and plant matter but we use the term ‘generalist’ for simplicity). *C. beltrani* is the most similar species in Chichancanab in both morphology and ecology to the coastal sister species, *C. artifrons* (Martin & Wainwright, 2011). We compared the first two major dimensions of intraspecific generalist morphological variance (first and second principal components of the *P* matrix) with the major dimensions of species divergence (first and second linear discriminant axes for 3 species) in each adaptive radiation. Similarly, we compared the first two major dimensions of intraspecific generalist dietary variance (first and second non-metric multidimensional scaling axes of intestinal contents) with the major dimension of species divergence in diet (canonical axis of the first principal coordinate axis) for each adaptive radiation. In nearly all cases, morphological and dietary variance within generalist populations was aligned with major axes of species divergence within each adaptive radiation.

## Methods

### Sampling

We collected Chichancanab pupfishes in July 2022 from the entrance to the lake at La Presumida in Quintana Roo, Mexico. Additional voucher specimens of these species are catalogued in the Museum of Vertebrate Zoology (MVZ:Fish:838,1410-1499, 1536) and the Universidad Nacional Autonoma de Mexico Ichthyology collections. We used a 6 m x 1.5 m seine net with 1/8” inch mesh size. Fish were euthanized immediately in the field in an overdose of buffered MS-222 following approved animal use protocols from the University of California, Berkeley IACUC (AUP-2021-07-14515-1 and AUP-2021-02-14062-1). Euthanized fish were immediately preserved in 100% ethanol. Ethanol was changed after 1-2 days for optimal preservation and specimens were stored at room temperature. Adult specimens for this study were collected randomly from a single collection site on a single day, resulting in one *C. simus* individual, 3 *C. labiosus*, and 105 *C. beltrani* in the final sample.

In July 2011, we sampled 12 hypersaline lakes on SSI for both generalist (*C. variegatus*) and specialist pupfishes (if present) and 10 hypersaline lakes on four neighboring islands (Cat, Long, Acklins, New Providence) containing *C. variegatus* populations as originally described in Martin (2016), resulting in a sample of 421 *C. variegatus* individuals. Individuals were collected in July, 2011 and additional sample details are described in Martin (2016). Voucher specimens from these collections are catalogued in the Museum of Vertebrate Zoology (MVZ:Fish:41:56, 501:513). Fish were sampled by seine or hand net and euthanized immediately in an overdose of buffered MS-222 following animal use protocols approved by the University of California, Davis Institutional Animal Care and Use Committee (IACUC protocols #15908 and #17455) and stored in 100% ethanol as described above.

### Morphometrics

Pupfish specimens from each adaptive radiation were measured in different labs using different external or skeletal measurement protocols for different sets of landmarks. This provides additional independence and robustness in our tests of ecological and morphological lines of least resistance across different morphometric and dietary datasets with independent investigators.

Chichancanab specimens were measured in the Basic Sciences laboratory at the Instituto Tecnológico de Felipe Carrillo Puerto. The following eleven external linear measurements were collected from each specimen using digital calipers: standard length (SL), caudal fin length (total length – SL), eye diameter, lower jaw length (tip of the most anterior tooth on the dentary to the approximate lower jaw joint just posterior to the ventral protruding tip of maxilla), snout length (tip of the most anterior tooth to the anterior edge of the orbit), interocular distance, head length (tip of the anterior tooth to the posterior edge of the operculum), head height (dorsal insertion of the epaxial muscle to the ventral edge of the suspensorium), preopercle height, body depth (first ray of dorsal fin to first ray of anal fin), and caudal peduncle height (minimum height anterior to the caudal fin). This set of external measurements was further described and diagrammed in Martin and Wainwright (2013).

Bahamian specimens were first cleared and double-stained with alizarin red and alcian blue following Dingerkus and Uhler (1977) before measurement of 28 linear skeletal traits using image analysis software. These measurements were originally described and published in Martin (2016). In brief, the skull of each specimen was photographed on both lateral sides with jaws adducted for a clear view of the jaw joint. Specimens were photographed and measured on both lateral sides and the mean was used to reduce measurement error. Thirty-two landmarks were digitized using tpsdig2 software (Rohlf 2001) and converted to 29 linear distances, defined and illustrated in Martin (2016): 1) lower jaw length, 2) dentary length, 3) lower jaw opening lever, 4) lower jaw closing lever, 5) angular coronoid width, 6) angular length, 7) retroarticular width, 8) tooth length, 9) premaxilla width, 10) premaxilla length, 11) upper jaw length, 12) premaxilla thickness, 13) dentigerous process width, 14) ascending process length, 15) maxilla length, 16) maxilla rotating arm, 17) maxilla thickness, 18) maxilla head protrusion, 19) orbit diameter, 20) neurocranium length, 21) neurocranium height, 22) palatine length, 23) head height, 24) pectoral height, 25) pectoral length, 26) adductor mandibulae (preopercle) height, 27) adductor mandibulae length, 28) quadrate length, 29) head size.

### Intestinal content analyses

*Cyprinodon* do not have stomachs, but only undifferentiated intestinal tracts (Day *et al*., 2011; Heras & Martin, 2022). Following exterior measurements and before clearing and staining of Bahamian specimens, the intestines were dissected from each individual and placed in sealed sample containers with 90-100% alcohol. All containers were labeled, indicating the name of the species and the tag number of the specimen to associate intestinal data with morphometrics.

Each intestinal tract was placed in a Petri dish and examined under a stereoscopic microscope. Contents were separated and identified to major taxonomic categories. For Chichancanab specimens, total surface areas of each dietary item were estimated under 10-50x magnification distributed over the surface of the Petri dish with a sheet of millimeter grid paper. Items were separated into seven categories: whole fish or skin/scales, bivalves, gastropods (including empty shells), arthropods, copepods, nematodes, plant matter (including macroalgae), and detritus. There are no polychaetes in Chichancanab and very rare dietary components (such as Odonata) were collapsed into larger taxonomic categories. There are no scale-eating specialists in Chichancanab, so individual scales were almost never observed and collapsed into the whole fish category.

For Bahamian specimens, total surface areas of each component were estimated under 10-50x magnification using a Sedgwick-rafter cell following Martin and Wainwright (2013c). Items were separated into ten categories: fish scales, bivalves, gastropods, arthropods, Odonata, ostracods, annelids, polychaetes, macroalgae, and angiosperm plant material. Macroalgae and angiosperms were separated for Bahamian dietary categories because these lakes are hypersaline whereas Laguna Chichancanab is slightly brackish and does not contain any macroalgae. Odonata were sufficiently distinctive within Bahamian gut contents to warrant their own category. We then calculated the proportional contribution of each item to the total intestinal contents for each individual, before calculating mean proportions for each population.

### Statistical analyses

We used linear discriminate analyses to calculate the multivariate dimensions of morphological species divergence within each adaptive radiation. Analyses used the lda function in the MASS package (Ripley *et al*., 2013) in R (R Core Team & Others, 2013). Chichancanab species were discriminated along a single axis of generalist versus trophic specialist (*n* = 4) due to the small sample size of specialists. For the proportional dietary data, we first converted values to a Bray-Curtis similar index. We then used canonical analysis of principal coordinates to calculate the multivariate dimensions of dietary species divergence within each adaptive radiation using the vegan package (Oksanen *et al*., 2013) in R.

To independently measure the major dimensions of intraspecific morphological diversity within Chichancanab and Bahamian generalist populations, we used principal component analysis of each size-corrected dataset using the princomp function in R. Size-correction was performed by taking the residuals from a linear regression of log-transformed trait measurements relative to log-transformed standard length (SL). To assess intraspecific dietary diversity, we used non-metric multidimensional scaling of the Bray-Curtis similarity indices for each dietary dataset using the metaMDS function in the vegan package (Oksanen *et al*., 2013). Importantly, these intraspecific analyses of major dimensions of variance were conducted only on generalist populations with all specialist species excluded.

Finally, to test the hypotheses of morphological and ecological lines of least resistance, we compared the two principal vectors of intraspecific diversity to the major discriminate axes of interspecific divergence within each adaptive radiation. For the morphological data, we compared the first two intraspecific principal components to the linear discriminate axes separating the species within each radiation using linear regression. For the dietary data, we compared the first two non-metric multidimensional scaling axes to the first canonical axis of principal coordinates using linear regression.

## Results

### Dietary specialization within Laguna Chichancanab radiation

Both radiations show the beginnings of dietary specialization in addition to retaining the ancestral omnivorous/detritivorus diet of generalist pupfishes *C. variegatus* and *C. beltrani*, similar to other young adaptive radiations (De León *et al*., 2014; Galvez *et al*., 2022). In Chichancanab, only one *simus* individual was collected in our random sample from the lake, but this individual was one out of only two fish in our sample of 108 that contained copepod remains in its intestines (Fig. 1). This species was originally described as a zooplanktivore based only on behavioral observations of its unique shoaling and zooplankton feeding behavior (Humphries & Miller, 1981; Plath & Strecker, 2008). The only subsequent dietary study of this species following the invasion of *Oreochromis niloticus x mossambicus* and *Astyanax cf. bacalarensis* reported only detritus in the gut contents (Horstkotte & Strecker, 2005; Schmitter-Soto, 2017). Although our sample is too small for formal statistics, this represents the first dietary record of zooplankton-feeding in this species (Humphries & Miller, 1981; Stevenson, 1992; Horstkotte & Strecker, 2005).

**Figure 1.**
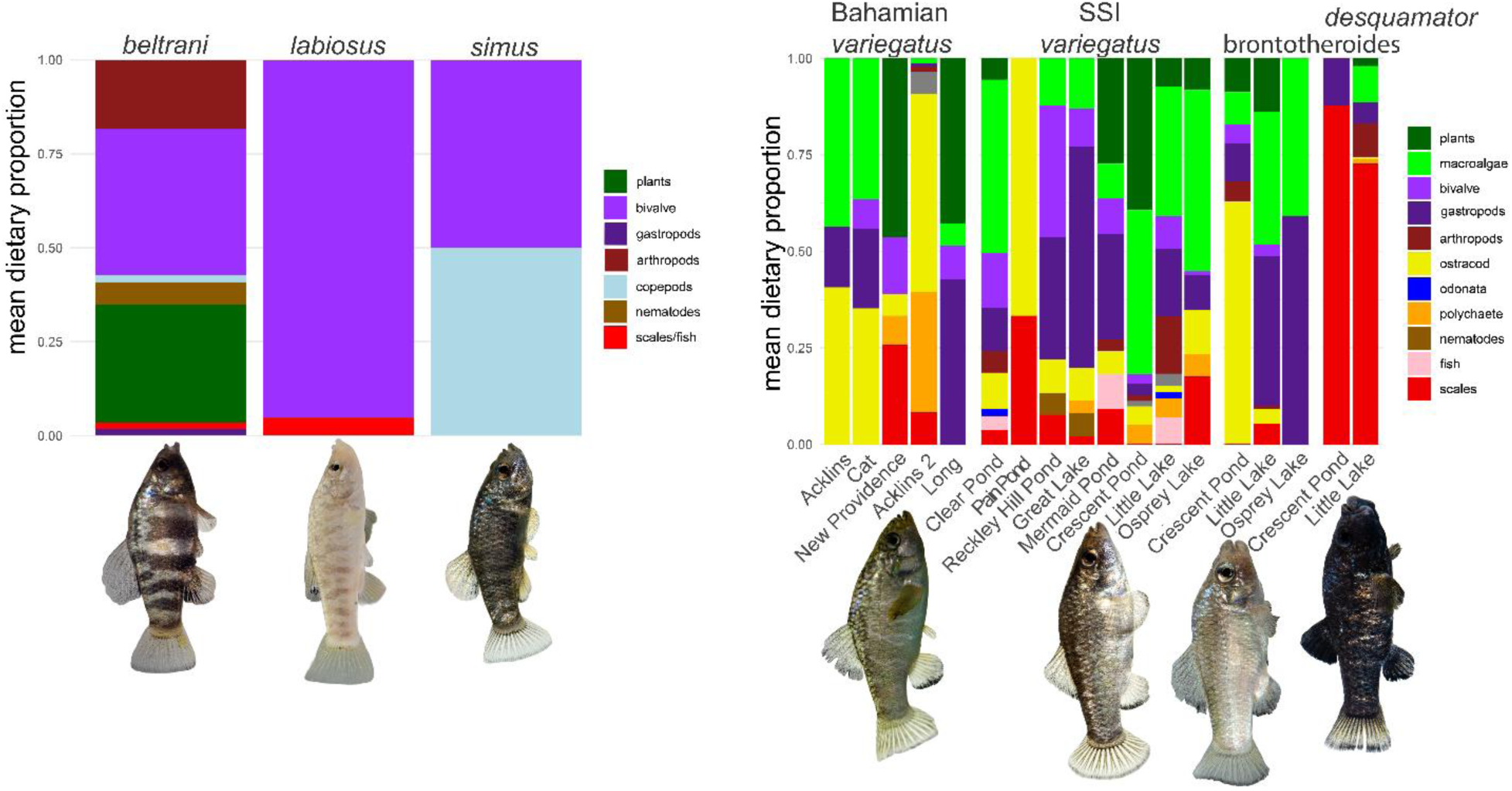
Dietary components in two adaptive radiations. Mean dietary proportions from surface area of intestinal contents for each species in a) Laguna Chichancanab (*C. beltrani, labiosus,* and *simus*) and b) SSI (*C. variegatus, brontotheroides, desquamator*) and *variegatus* populations from four neighboring Bahamian islands (Cat, New Providence, Long, and Acklins). Representative photos of each species are shown from captive-bred common garden populations except for *C. beltrani* and *C. labiosus*. Allopatric *C. variegatus* populations on SSI are Clear Pond, Pain Pond, and Reckley Hill Pond.

Similarly, *C. labiosus*, originally described as feeding on snails and “small crustaceans” (Humphries & Miller, 1981), was later considered an amphipod specialist based on the over-representation of this dietary item in its gut contents relative to other species (Horstkotte & Strecker, 2005). In the three *C. labiosus* within our random sample, we detected no amphipod (or any arthropod) remains despite the abundance of arthropods consumed by *C. beltrani* (Fig. 1). However, all three *C. labiosus* were among the top five consumers of bivalves (clams) in our sample of 108 fish (Fig. 1: purple bars). Although our sample is too small for formal statistics, this represents the first dietary record of bivalve feeding in this species and begs the question of whether their species-characteristic enlarged fleshy lips might provide an adaptation for this resource, as already speculated by Humphries & Miller (1981). In contrast to specialists from SSI, zero plant material was observed in the guts of Chichancanab specialists.

### Dietary specialization within San Salvador Island radiation and neighboring islands

On SSI, both lake populations of *C. desquamator* showed a clear abundance of fish scales in their diet from 75% in Little Lake to 87% in Crescent Pond which also was the only population with no plant or macroalgae material in its diet (Fig. 1). Both scale-eaters also consumed gastropods and arthropods. The molluscivore *C. brontotheroides* consumed a comparable amount of plant/macroalgae and an excess of gastropods in the Little/Osprey Lake populations relative to generalist *C. variegatus*. In Crescent Pond, *C. brontotheroides* consumed a substantial excess of ostracods relative to *C. variegatus* except for one *C. variegatus* population in Pain Pond with a lower sample size (Fig. 1). Finally, there were no clear differences among *C. variegatus* populations on SSI versus neighboring Bahamian islands or *C. variegatus* populations in sympatry or allopatry with specialists on SSI; all consumed a mix of plant material, gastropods, polychaetes, and arthropods such as ostracods, amphipods, and Odonata (dragonfly nymphs), plus occasionally whole pupfish.

### Morphological and dietary divergence within each adaptive radiation

Both adaptive radiations showed significant evidence of species divergence in size-corrected morphology and diet (Fig. 2). Discriminant analysis based on external landmarks separated trophic specialist individuals in the Chichancanab radiation (*C*. *simus + C*. *labiosus*: *n* = 4) from *beltrani* generalists (n = 98) on LD1 with a classification accuracy of 97% (Fig. 2a). The strongest loadings on this axis were orbit diameter and head length, consistent with the elongated snout of *C*. *labiosus* and enlarged eyes of both *C*. *labiosus* and *C. simus*. Among Bahamian pupfishes, discriminant analysis separated *desquamator* from other species with 100% accuracy (*n* = 23) and *C*. *brontotheroides* from other species with 92.3% accuracy (*n* = 13; Fig. 2b). The strongest loadings on LD1 separating *C*. *desquamator* from other species were dentary length and premaxilla length, consistent with the elongated oral jaws of *C*. *desquamator*. The strongest loadings on LD2 separating *C*. *brontotheroides* from other species were dentary length (negative loading) and maxillary head length, consistent with the protruding maxillary head and short oral jaws of *C*. *brontotheroides*.

**Figure 2.**
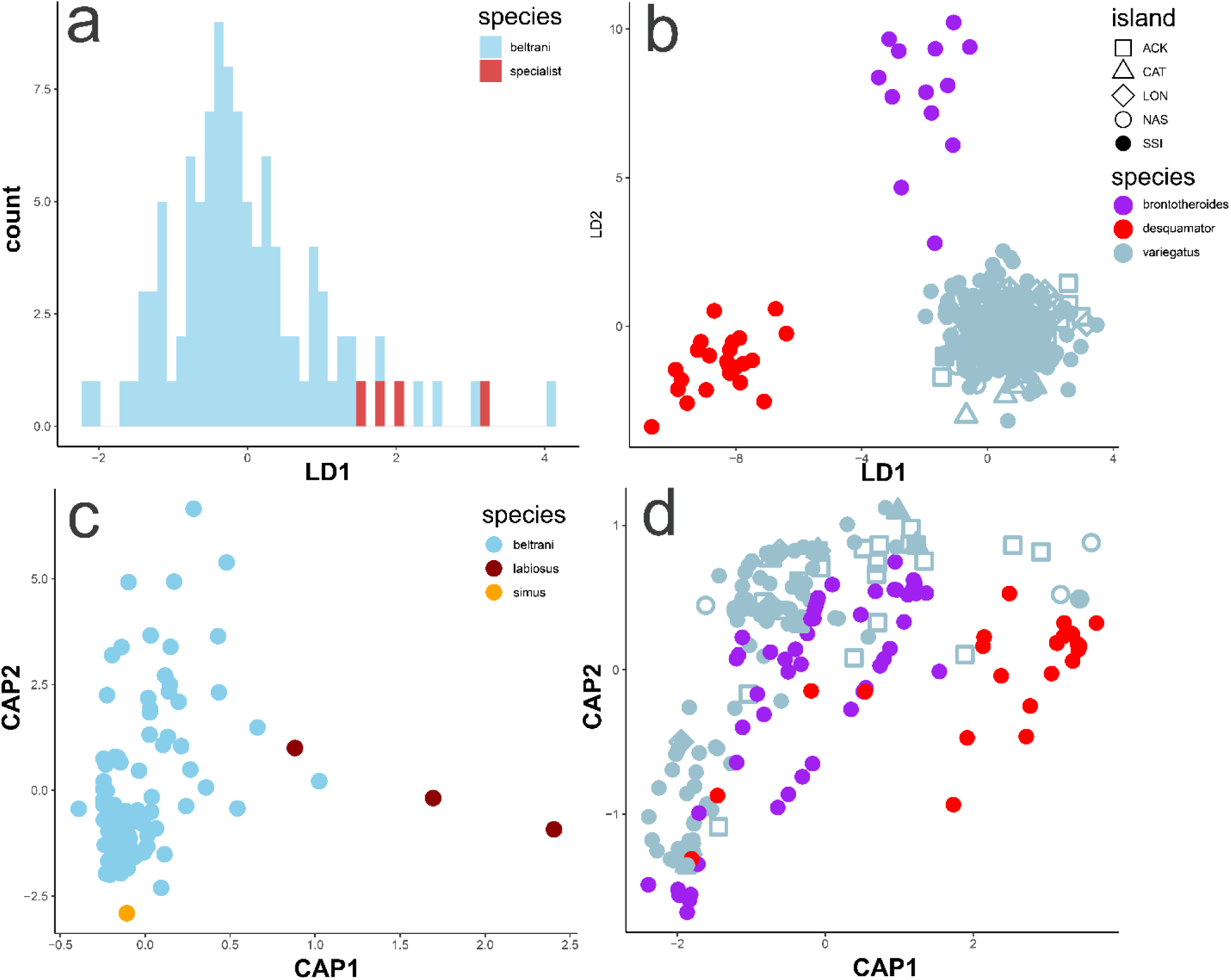
Multivariate discriminate analyses of morphology and diet in two adaptive radiations. Chichancanab (first column) and San Salvador Island (second column) pupfish species flocks show divergence in craniofacial morphology (first row) and diet from quantification of intestinal contents (second row). a) Four trophic specialist individuals (3 *C*. *labiosus* and 1 *C*. *beltrani*) from Chichancanab show substantial separation along linear discriminate axis 1 (LD1) relative to generalist individuals in blue (*C*. *beltrani*). Specialists were pooled into a single specialist vs. generalist category due to low sample size, resulting in only a single discriminant axis. b) Both trophic specialist species from SSI (SSI) show complete separation along linear discriminate axis 1 (LD1) for the scale-eater *C*. *desquamator* and along linear discriminate axis 2 (LD2) for the molluscivore *C*. *brontotheroides* relative to generalist populations (*C*. *variegatus*) from SSI (blue filled circles) and neighboring Bahamian islands (open shapes). c) The bivalve-specialist *C*. *labiosus* shows significant separation in diet along the canonical axis of principal coordinate 1 (CAP1) relative to generalist individuals in blue (*C*. *beltrani*). d) Both trophic specialist species from SSI (SSI; *C*. *desquamator* and *C*. *brontotheroides*) show significant separation along the canonical axis of principal coordinate 1 (CAP1) relative to generalist populations (*C. variegatus*) from SSI (blue filled circles) and neighboring Bahamian islands (open shapes) (Acklins, Cat, Long, and Nassau [New Providence] Islands).

Similarly, Chichancanab showed strong separation between predator and detritivore diets along the first canonical axis of principal coordinates analysis (CAP1), which provides a discriminant analysis of the categorical dietary data (Fig. 2c). Both SSI specialists *C*. *desquamator* and *C*. *brontotheroides* also showed substantial dietary separation along CAP1 (Fig. 2d). At the same time, we observed minimal separation between SSI *C*. *variegatus* and neighboring *C*. *variegatus* populations by diet (Fig. 2d).

### Intraspecific morphological and dietary variance within generalist populations

The major axis of phenotypic variance within *C*. *beltrani* in the Chichancanab radiation (PC1) explained 22.7% of the overall external morphological variation measured with primary loadings of 1) head height and 2) opercle length. The second major axis of phenotypic variance (PC2) explained 18% of overall variation with major loadings of 1) body depth and 2) caudal peduncle height (Fig. 3a). Similarly, the major axis of phenotypic variance across Bahamian *C*. *variegatus* generalist populations, including SSI, explained 35.9% of the overall craniofacial skeletal variation, including major loadings of 1) suspensorium length and 2) oral jaw length. The second major axis of phenotypic variance (PC2) explained 7.9% of overall variation with major loadings of 1) orbit diameter and 2) head length (Fig. 3b). There was no clear separation between SSI variegatus and neighboring islands, although certain island populations, such as Long Island, did cluster in smaller regions of the morphospace relative to the SSI generalist populations perhaps due to the greater number of lake populations sampled from SSI.

**Figure 3.**
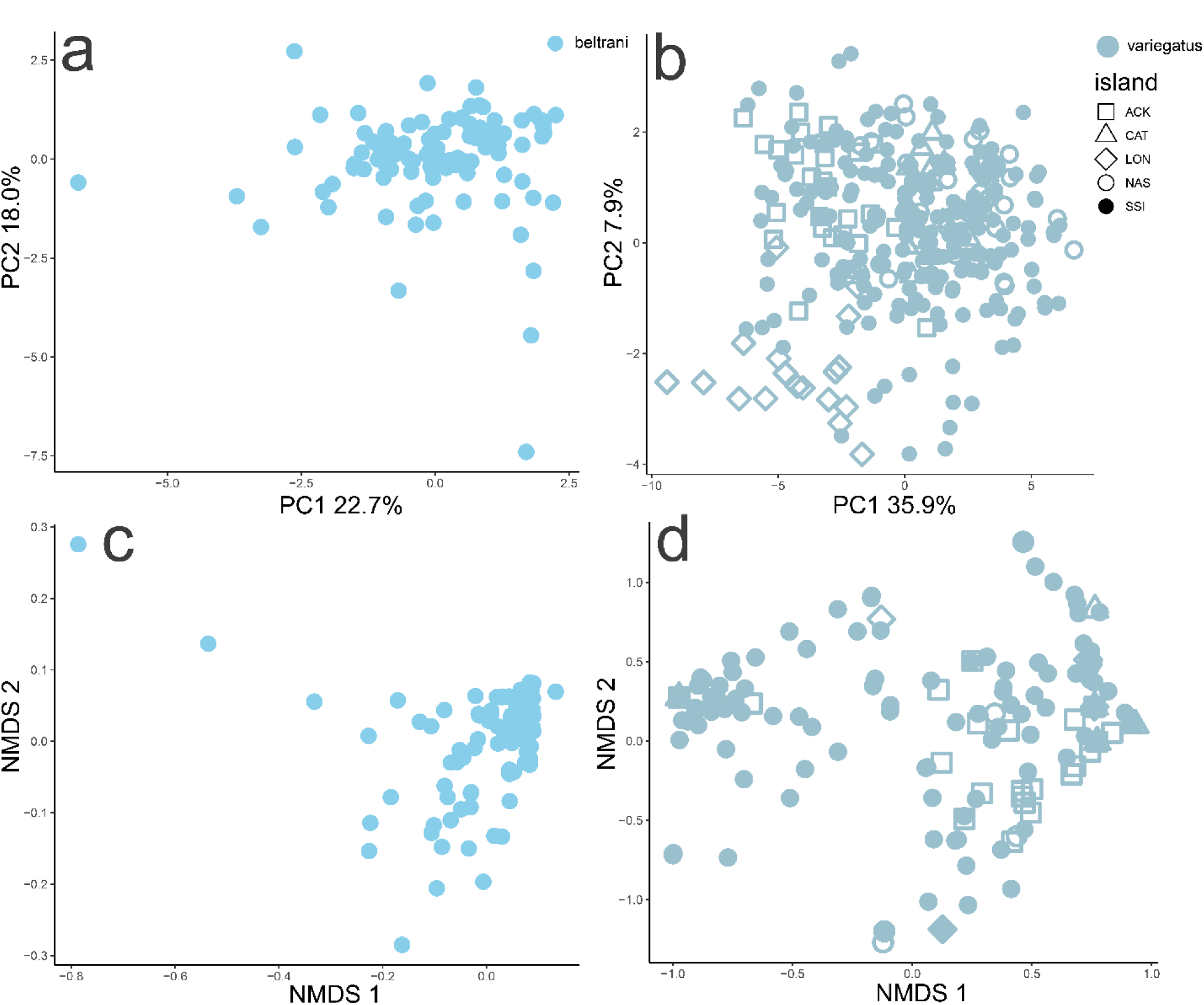
Major axes of intraspecific craniofacial and dietary variation for generalist populations within two adaptive radiations including neighboring Bahamian islands. Chichancanab (first column) and Bahamian (second column) generalist populations show substantial intraspecific variation in craniofacial morphology (first row) and diet from quantification of intestinal contents (second row). First two principal components of trophic morphology for a) nine external linear measurements of wild-collected *C*. *beltrani* from Chichancanab, Mexico and b) 24 linear measurements of cleared and alizarin-stained wild-collected specimens of *C*. *variegatus* from SSI (filled circles) and four neighboring Bahamian islands (Acklins, Cat, Long, and New Providence Islands). First two principal coordinate axes of diet (NMDS1-2: non-metric dimensional scaling) from intestinal content analyses of c) wild-collected *C. beltrani* from Chichancanab, Mexico and d) wild-collected specimens of *C. variegatus* from SSI (filled circles) and four neighboring Bahamian islands (Acklins, Cat, Long, and New Providence Islands).

Similarly, non-metric multidimensional scaling plots indicated clear individual variation in diet within each radiation and no clear separation between the diets of SSI generalists and neighboring Bahamian populations (Fig. 3c-d).

### Adaptive radiation along morphological lines of least resistance

Both adaptive radiations showed significant associations between major dimensions of intraspecific morphological variance and interspecific divergence during adaptive radiation. In Chichancanab, there was a highly significant association between the first major axis of morphological diversity (PC1) within the *C*. *beltrani* generalist population and the discriminant axis (LD1) separating this species from trophic specialists (linear regression, *P* < 10e^-16^; Fig. 4a). Intraspecific generalist diversity explained 55% of the variance in species divergence within this radiation, despite being calculated independently without the influence of trophic specialist morphology. There was no significant relationship between discriminant axis 1 (LD1) and the second major axis of morphological variation in *C*. *beltrani* generalists (PC2; linear regression, P = 0.883; Fig. 4b).

**Figure 4.**
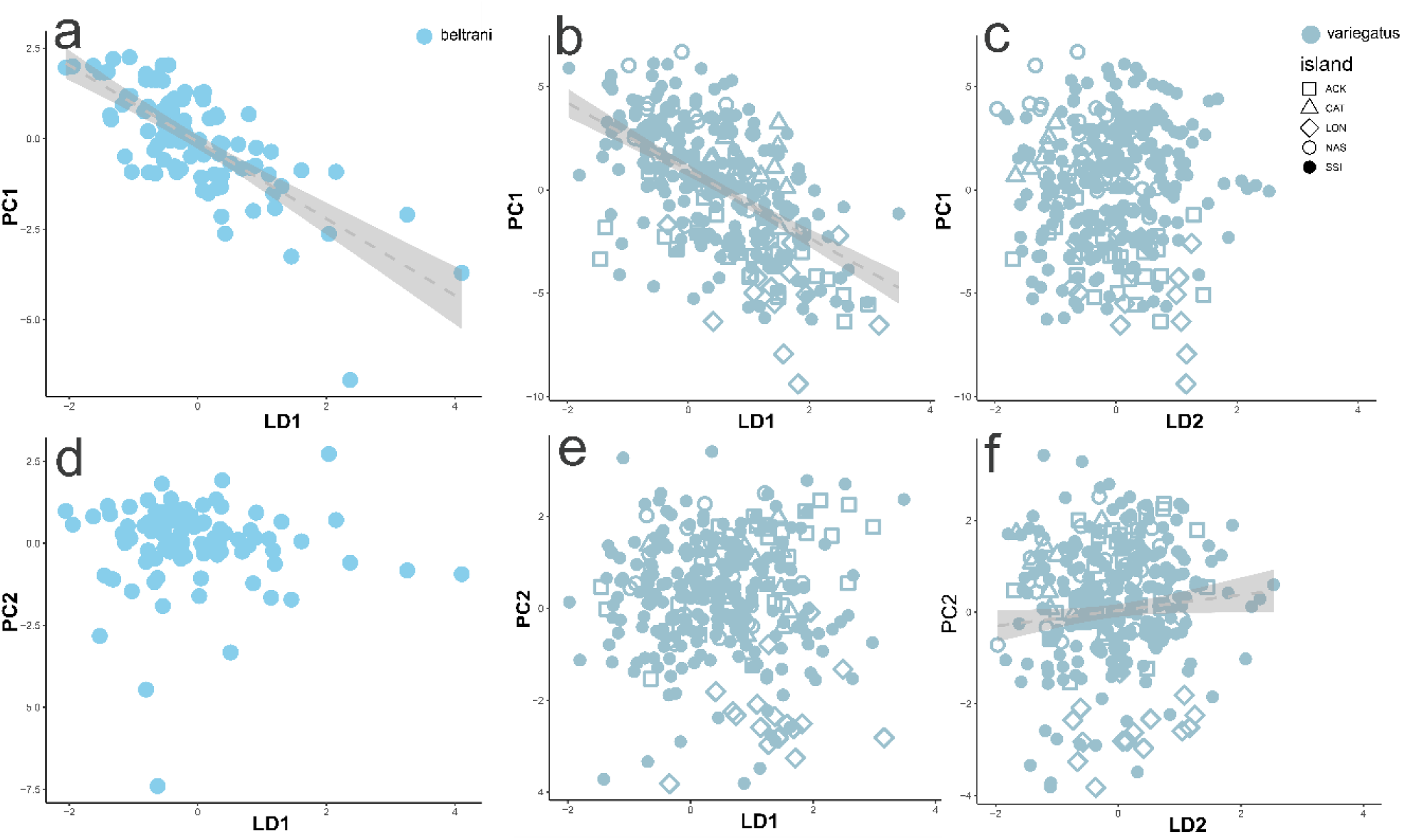
Major axes of intraspecific generalist craniofacial variation were significantly associated with discriminate axes of craniofacial divergence within two adaptive radiations. Laguna Chichancanab (first column) and Bahamian (second and third columns) generalist pupfish populations show strong alignment between their first principal component axis of variation in trophic morphology (PC1, Fig. 3) and primary discriminate axis (LD1) separating trophic specialists from generalists in each radiation (Fig. 2). a,c) The primary axis of *C. beltrani* morphological diversity (PC1) aligns with generalist-specialist morphological divergence (LD1) within the Chichancanab radiation, Mexico. b,d) The primary and secondary axes of *C. variegatus* craniofacial diversity (PC1-2) from SSI (blue filled circles) and neighboring Bahamian islands (open shapes) (Acklins, Cat, Long, and New Providence Islands) aligns with generalist-specialist craniofacial divergence along LD1 (discriminating the scale-eater) and LD2 (discriminating the molluscivore) within the SSI radiation. Note that principal component analyses were calculated only within generalist populations (Fig. 3) independent of specialists to prevent bias.

In the SSI radiation, there was a highly significant association between the first major axis of morphological diversity (PC1) within Bahamian *C*. *variegatus* generalist populations and the discriminant axis (LD1) separating this species from *C*. *desquamator* (linear regression, *P* < 2e^-16^; Fig. 4b). Intraspecific generalist diversity on PC1 explained 29.6% of the variance in species divergence within the SSI radiation on the primary discriminant axis. There was no significant relationship between discriminant axis 2 (LD2) and the second major axis of morphological variation in *C*. *variegatus* generalists (PC1; linear regression, *P* = 0.480; Fig. 4c) nor between discriminant axis 1 and PC2 (*P* = 0.480; Fig. 4e). However, there was a significant, albeit weaker, association between the second major axis of morphological diversity (PC2) within Bahamian *C*. *variegatus* generalist populations and the second discriminant axis (LD2) separating *C*. *brontotheroides* from other species (linear regression, *P* = 0.019; Fig. 4f). Intraspecific generalist diversity on PC2 explained 1.2% of the variance in species divergence within the SSI radiation on the second discriminant axis (Fig. 4f).

### Adaptive radiation along ecological lines of least resistance

Both adaptive radiations showed significant associations between the major discriminant axis of dietary divergence during radiation (CAP1) and both primary and secondary dimensions of intraspecific variance in diet among generalists (NMDS1-2; Fig. 5). In Chichancanab, there was a significant association between the canonical axis of principal coordinate 1 separating specialist diets from generalists and both the first and second major axes of dietary variance (NMDS1-2) in the *beltrani* generalist population (NMDS1: linear regression, *P* = 0.027; Fig. 5a; NMDS2: linear regression, *P* = 0.027, Fig 5c). Intraspecific generalist dietary diversity on NMDS1 explained 3.9% and NMDS2 explained 3.9% of species divergence in diet within the Chichancanab radiation.

**Figure 5.**
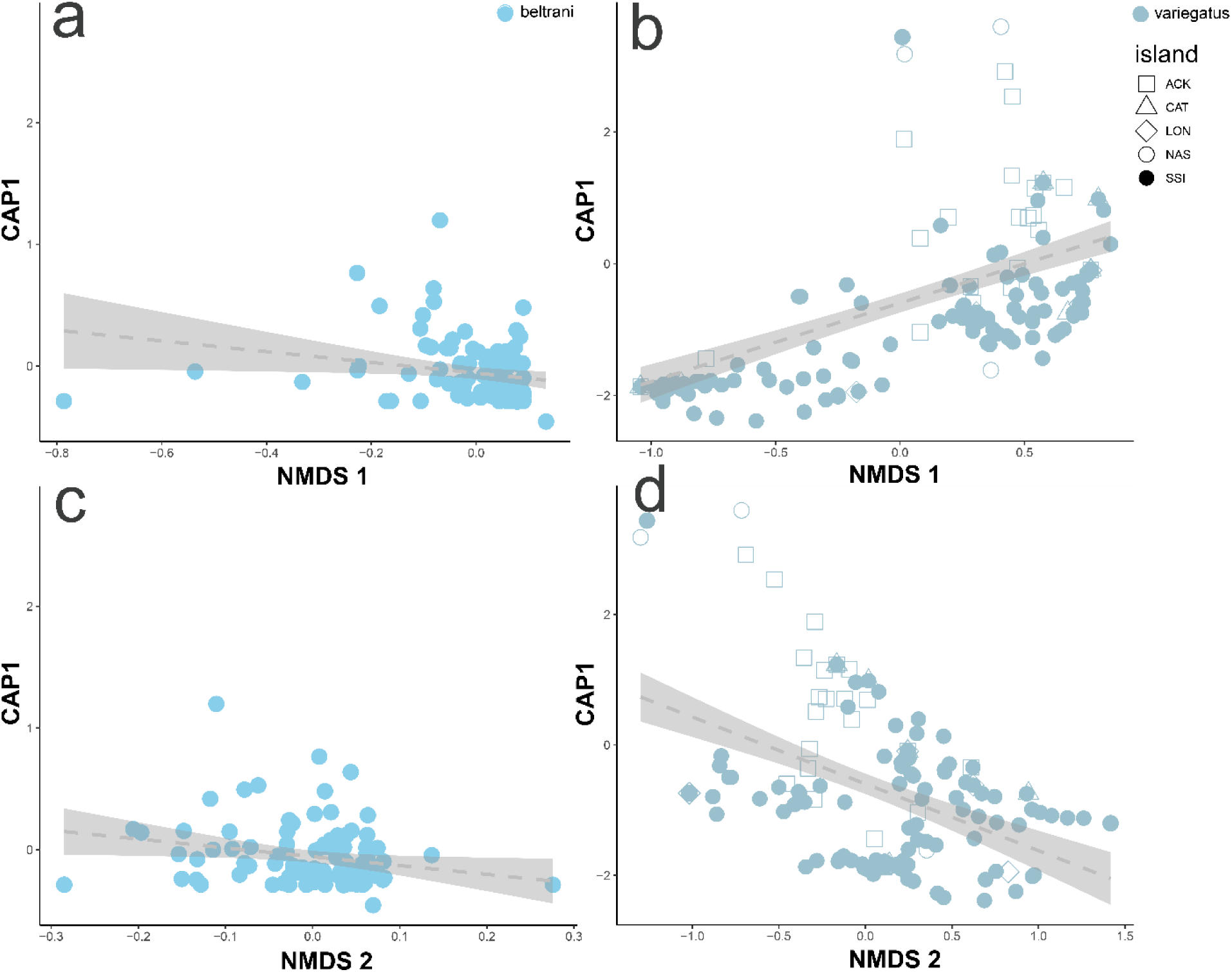
Major axes of intraspecific generalist dietary variation were significantly associated with the major axis of interspecific dietary variation within two adaptive radiations. Chichancanab (first column) and Bahamian (second column) generalist pupfish populations show strong alignment between their first two principal coordinate axes of diet (NMDS1-2: non-metric dimensional scaling; Fig. 3) and the canonical axis of principal coordinate 1 (CAP1; Fig. 2) separating trophic specialists from generalists in each radiation. a,c) *C. beltrani* dietary diversity aligns with generalist-specialist dietary divergence along CAP1 within the Chichancanab radiation, Mexico. b,d) *C. variegatus* dietary diversity from SSI (blue filled circles) and neighboring Bahamian islands (open shapes) (Acklins, Cat, Long, and New Providence Islands) aligns with generalist-specialist dietary divergence along CAP1 within the SSI radiation. Note that NMDS principal coordinate axes were calculated only within generalist populations (Fig. 3) independent of specialists to prevent bias.

In the SSI radiation, there was a highly significant association between the canonical axis of principal coordinate 1 separating specialist diets from generalists and both the first and second major axes of dietary variance (NMDS1-2) in Bahamian *C*. *variegatus* generalist populations (NMDS1: linear regression, *P* = 10e^-16^; Fig. 5b; NMDS2: linear regression, *P* = 10e^-13^, Fig 5d). Intraspecific generalist dietary diversity on NMDS1 explained 33% and NMDS2 explained 20.3% of the variance in species divergence within the SSI radiation.

## Discussion

Here we found nearly universal congruence between major morphological and dietary “lines of least resistance”, such that the multivariate dimensions representing the greatest variance within generalist populations aligned with the multivariate directions of species divergence within two independent adaptive radiations of *Cyprinodon* pupfishes geographically separated across the Caribbean. The major dimension of craniofacial variance in outgroup generalist populations was significantly associated with the primary axis of species divergence in both radiations (Fig. 4a-b). The second largest dimension of craniofacial variance in Bahamian generalist populations was also associated with the secondary axis of species divergence (separating the molluscivore specialist) on SSI (Fig. 4f). Moreover, both the first and second largest dimensions of dietary variance in generalist populations were significantly associated with the primary canonical axis discriminating the diets of specialists from generalists within each radiation (Fig. 5). Overall, this suggests that adaptive radiations tend to leverage major ecological and morphological dimensions of variance that pre-exist in their founding population (Schluter, 1996, 2024; Marroig & Cheverud, 2005; McGlothlin *et al*., 2018; Chan *et al*., 2024). This contradicts the widespread view that ecological novelty in new environments, not pre-existing dimensions of variance, is the primary driver of the direction of species divergence during radiation (Yoder *et al*., 2010; Gillespie *et al*., 2020; Stroud *et al*., 2025).

These results are particularly striking given that we also clearly document novel resource specialization and trophic innovation within each radiation (Martin & Wainwright, 2011). The conservation status of the Chichancanab specialists means that, despite limited sampling in this study, we were able to place critical ecological and morphological observations within the context of the larger radiation, an opportunity that is likely lost for future investigators. A different set of trophic specialists is unique to SSI: up to 87% of the diet of *C*. *desquamator* was scales with minimal plant matter unlike all other pupfishes; an excess of gastropods or ostracods was observed in the diet of *C*. *brontotheroides* across three lake populations (Fig. 1; also see (Martin & Wainwright, 2013c).

One interpretation is that the outgroup Bahamian *C*. *variegatus* populations and *C*. *beltrani* population in Chichancanab still contain additive genetic variation for adaptive craniofacial traits and correlated dietary preferences for these novel trophic resources which was co-opted for species divergence after colonization of these largely predator-free habitats (e.g. see (Beltman, Haccou, & ten Cate, 2004; Gavrilets & Vose, 2005; Ricklefs, 2010; Tanentzap *et al*., 2015; Basnou *et al*., 2016)). Indeed, all generalist populations were observed to feed rarely on the same resources as specialists, including copepods, bivalves, fish scales, gastropods, and ostracods. Alternatively, similar ecological opportunities across Caribbean lake populations may be shaping both generalist populations and the adaptive radiations in parallel, creating an apparent association between generalist dimensions of variance and axes of species divergence during adaptive radiation, but no underlying mechanism of adaptive divergence facilitated by pre-existing variation within the founding populations (e.g. see (Schluter *et al*., 2004; Kaeuffer *et al*., 2012b; Bailey, Rodrigue, & Kassen, 2015; Esquerré & Scott Keogh, 2016; McGirr & Martin, 2018; De Lisle, Bolnick, & Stuart, 2024)). Separating these alternative hypotheses is difficult, but fundamental to understanding whether lines of least resistance within the founding population shape the direction of subsequent species divergence during adaptive radiation or whether these congruent lines of least resistance simply reflect shared environmental selective pressures among similar habitats that shift over time for both generalist populations and radiating species. If similar environments drive parallels, we might expect environmental similarity to best predict congruence between generalist diversity and species divergence, whereas phylogenetic or geographic distance would be the best predictors if pre-existing lines of least resistance shape radiation. In support of the latter hypothesis, there appears to be no difference in major dimensions of craniofacial or dietary variance between neighboring Bahamian populations and SSI generalists, suggesting that these dimensions are quite stable across environments and over phylogenetic distances vastly longer than the recent origins of both radiations 8 ka to 15 ka (Hodell, Brenner, & Curtis, 2005; Strecker, 2006a; Richards *et al*., 2021).

One major caveat is that intraspecific dimensions of ecological and morphological variance did not explain the majority of species divergence in any case, and sometimes explained a very small percentage to total variation despite significance, such as Chichancanab divergence in diet. Thus, these adaptive radiations still proceeded largely in unpredictable directions relative to the putative major dimensions of ecological and morphological variance in the founding populations. However, our study suggests that pre-existing lines of least ecological resistance provide some predictive power and may frequently shape or constrain directions of adaptive divergence in nature.

We believe this study provides a new framework for assessing ecological lines of least resistance in addition to the burgeoning literature on genetic lines of least resistance originating from Schluter’s original hypothesis (Schluter, 1996). We found surprising support for this hypothesis and the more traditional prediction that dimensions of the *P* matrix constrain adaptive radiation, suggesting that adaptive divergence was somewhat channeled along existing dimensions of variation within the founding population. Importantly, we show this to be the case for two independent *Cyprinodon* radiations that diverged in different directions toward unique trophic specializations. We also provide critical natural history data on the diets of these species that may now be extinct in the wild.

## Acknowledgments

This research was funded by NSF CAREER 1749764, NIH 5R01DE027052-02, Daphne and Ted Pengelley Award from the Center for Population Biology, and ARCS Foundation awards to CHM. We thank the Gerace Research Center and Troy Day for logistical support in the field, and the governments of Mexico and the Bahamas BEST commission for permission to collect and export samples (Permit No. PPF/DGOPA-001/20). All research procedures and animal care protocols (AUP-2021-02-14062-1 and AUP-2021-07-14515) were approved by the University of California, Berkeley Animal Care and Use committee and the UC Davis Institutional Animal Care and Use committee. Claude.ai was used for assistance with R coding but otherwise AI was not used for any text generation or revision.

## Data Accessibility

All data will be uploaded to the Dryad digital repository upon acceptance.

## References

Arnold SJ, Bürger R, Hohenlohe PA, Ajie BC & Jones AG. 2008. Understanding the evolution and stability of the G-matrix. Evolution; international journal of organic evolution 62: 2451–2461.

Baguette M, Bertrand JAM, Stevens VM & Schatz B. 2020. Why are there so many bee-orchid species? Adaptive radiation by intra-specific competition for mnesic pollinators. Biological reviews of the Cambridge Philosophical Society 95: 1630–1663.

Bailey SF, Rodrigue N & Kassen R. 2015. The effect of selection environment on the probability of parallel evolution. Molecular biology and evolution 32: 1436–1448.

Barton N & Turelli M. 1987. Adaptive landscapes, genetic distance and the evolution of quantitative characters. Genetical research 49: 157–173.

Basnou C, Vicente P, Espelta JM & Pino J. 2016. Of niche differentiation, dispersal ability and historical legacies: what drives woody community assembly in recent Mediterranean forests? Oikos (Copenhagen, Denmark) 125: 107–116.

Beausoleil MO, Carrión PL, Podos J, Camacho C, Rabadán-González J, Richard R, Lalla K, Raeymaekers JAM, Knutie SA, De León LF, Chaves JA, Clayton DH, Koop JAH, Sharpe DMT, Gotanda KM, Huber SK, Barrett RDH & Hendry AP. 2023. The fitness landscape of a community of Darwin’s finches. Evolution; international journal of organic evolution 77: 2533–2546.

Beltman JB, Haccou P & ten Cate C. 2004. Learning and colonization of new niches: a first step toward speciation. Evolution; international journal of organic evolution 58: 35–46.

Berner D, Stutz WE & Bolnick DI. 2010. Foraging trait (co) variances in stickleback evolve deterministically and do not predict trajectories of adaptive diversification. Evolution.

Bolnick. 2004. Can intraspecific competition drive disruptive selection? An experimental test in natural populations of sticklebacks. Evolution; international journal of organic evolution 58: 608–618.

Bolnick DI, Svanbäck R, Araújo MS & Persson L. 2007. Comparative support for the niche variation hypothesis that more generalized populations also are more heterogeneous. Proceedings of the National Academy of Sciences of the United States of America 104: 10075– 10079.

Bolnick DI & Lau OL. 2008. Predictable patterns of disruptive selection in stickleback in postglacial lakes. The American naturalist 172: 1–11.

Bolnick DI & Paull JS. 2009. Morphological and dietary differences between individuals are weakly but positively correlated within a population of threespine stickleback. Evolutionary Ecology Research 11: 1217–1233.

Chan H, Colaco E, Martin CH & Evans KM. 2024. Adaptive radiation despite conserved modularity patterns in San Salvador Island Cyprinodon pupfishes and their hybrids. Evolutionary Journal of the Linnean Society 3: kzae013.

Day RD, German DP, Manjakasy JM, Farr I, Hansen MJ & Tibbetts IR. 2011. Enzymatic digestion in stomachless fishes: how a simple gut accommodates both herbivory and carnivory. Journal of comparative physiology. B, Biochemical, systemic, and environmental physiology 181: 603–613.

De León LF, Podos J, Gardezi T, Herrel A & Hendry AP. 2014. Darwin’s finches and their diet niches: the sympatric coexistence of imperfect generalists. Journal of evolutionary biology 27: 1093–1104.

De Lisle SP, Bolnick DI & Stuart YE. 2024. Predictable and divergent change in the multivariate P matrix during parallel adaptation. The American naturalist 204: 15–29.

De-Kayne R, Schley R, Barth JMI, Campillo LC, Chaparro-Pedraza C, Joshi J, Salzburger W, Van Bocxlaer B, Cotoras DD, Fruciano C, Geneva AJ, Gillespie R, Heras J, Koblmüller S, Matthews B, Onstein RE, Seehausen O, Singh P, Svensson EI, Salazar-Valenzuela D, Vanhove MPM, Wogan GOU, Yamaguchi R, Yoder AD & Cerca J. 2024. Why Do Some Lineages Radiate While Others Do Not? Perspectives for Future Research on Adaptive Radiations. : a041448.

Deng J, Cordero OX, Fukami T, Levin SA, Pringle RM, Solé R & Saavedra S. 2024. The development of ecological systems along paths of least resistance. Current biology: CB 34: 4813–4823.e14.

Dieckmann U & Doebeli M. 1999. On the origin of species by sympatric speciation. Nature 400: 354–357.

Dunker JC, St John ME & Martin CH. 2024. Phenotypic covariation predicts diversification in an adaptive radiation of pupfishes. Ecology and evolution 14: e11642.

Eroukhmanoff F, Eroukhmanoff F & Svensson E. 2011. Evolution and stability of the G- matrix during the colonization of a novel environment. Journal of evolutionary biology 24.

Esquerré D & Scott Keogh J. 2016. Parallel selective pressures drive convergent diversification of phenotypes in pythons and boas. Ecology letters 19: 800–809.

Evans KM, Larouche O, Watson SJ, Farina S, Habegger ML & Friedman M. 2021. Integration drives rapid phenotypic evolution in flatfishes. Proceedings of the National Academy of Sciences of the United States of America 118.

Fuselier L. 2001. Impacts of Oreochromis mossambicus (Perciformes: Cichlidae) upon habitat segregation among cyprinodontids (Cyprinodontiformes) of a species flock in Mexico. Revista de biologia tropical 49: 647–655.

Galvez JR, St John ME, McLean K, Touokong CD, Gonwouo LN & Martin CH. 2022. Trophic specialization on unique resources despite limited niche divergence in a celebrated example of sympatric speciation. Ecology of freshwater fish 31: 675–692.

Gavrilets S & Vose A. 2005. Dynamic patterns of adaptive radiation. Proceedings of the National Academy of Sciences of the United States of America 102: 18040–18045.

Gillespie RG, Bennett GM, De Meester L, Feder JL, Fleischer RC, Harmon LJ, Hendry AP, Knope ML, Mallet J, Martin C, Parent CE, Patton AH, Pfennig KS, Rubinoff D, Schluter D, Seehausen O, Shaw KL, Stacy E, Stervander M, Stroud JT, Wagner C & Wogan GOU. 2020. Comparing Adaptive Radiations Across Space, Time, and Taxa. The Journal of heredity 111: 1–20.

Gracida-Juárez CA, Schmitter-Soto JJ & Genner MJ. 2024. Community structure of indigenous fishes relative to habitat variation and invasive tilapia in lakes of Quintana Roo, Mexico. Environmental biology of fishes.

Heras J & Martin CH. 2022. Minimal overall divergence of the gut microbiome in an adaptive radiation of Cyprinodon pupfishes despite potential adaptive enrichment for scale-eating. PloS one 17: e0273177.

Hernandez LP, Adriaens D, Martin CH, Wainwright PC, Masschaele B & Dierick M. 2018. Building trophic specializations that result in substantial niche partitioning within a young adaptive radiation. Journal of anatomy 232: 173–185.

Hodell DA, Brenner M & Curtis JH. 2005. Terminal Classic drought in the northern Maya lowlands inferred from multiple sediment cores in Lake Chichancanab (Mexico). Quaternary science reviews 24: 1413–1427.

Horstkotte J & Strecker U. 2005. Trophic differentiation in the phylogenetically young Cyprinodon species flock (Cyprinodontidae, Teleostei) from Laguna Chichancanab (Mexico): TROPHIC DIFFERENTIATION IN A CYPRINODON SPECIES FLOCK. Biological journal of the Linnean Society. Linnean Society of London 85: 125–134.

Humphries JM. 1984. Cyprinodon verecundus, n. sp., a Fifth Species of Pupfish from Laguna Chichancanab. Copeia 1984: 58–68.

Humphries J & Miller RR. 1981. A remarkable species flock of pupfishes, genus Cyprinodon, from Yucatán, México@@@A remarkable species flock of pupfishes, genus Cyprinodon, from Yucatan, Mexico. Copeia 1981: 52.

Kaeuffer R, Peichel CL, Bolnick DI & Hendry AP. 2012a. Parallel and nonparallel aspects of ecological, phenotypic, and genetic divergence across replicate population pairs of lake and stream stickleback. Evolution.

Kaeuffer R, Peichel CL, Bolnick DI & Hendry AP. 2012b. Parallel and nonparallel aspects of ecological, phenotypic, and genetic divergence across replicate population pairs of lake and stream stickleback: Parallel and non-parallel evolution. Evolution; international journal of organic evolution 66: 402–418.

Knudsen R, Primicerio R, Amundsen PA & Klemetsen A. 2010. Temporal stability of individual feeding specialization may promote speciation. The journal of animal ecology 79: 161–168.

Marroig G & Cheverud J. 2005. Size as a line of least evolutionary resistance: Diet and adaptive morphological radiation in New World monkeys. Evolution 59: 1128–1142.

Martin CH. 2012. Weak disruptive selection and incomplete phenotypic divergence in two classic examples of sympatric speciation: cameroon crater lake cichlids. The American naturalist 180: E90–E109.

Martin CH & Feinstein LC. 2014. Novel trophic niches drive variable progress towards ecological speciation within an adaptive radiation of pupfishes. Molecular ecology 23: 1846– 1862.

Martin CH & Richards EJ. 2019. The paradox behind the pattern of rapid adaptive radiation: how can the speciation process sustain itself through an early burst? Annual review of ecology, evolution, and systematics 50: 569–593.

Martin & Wainwright PC. 2011. Trophic novelty is linked to exceptional rates of morphological diversification in two adaptive radiations of Cyprinodon pupfish: Cyprinodon adaptive radiations. Evolution; international journal of organic evolution 65: 2197–2212.

Martin & Wainwright PC. 2013a. A remarkable species flock of Cyprinodon pupfishes endemic to San Salvador Island, Bahamas. Bulletin of the Peabody Museum of Natural.

Martin CH & Wainwright PC. 2013b. Multiple fitness peaks on the adaptive landscape drive adaptive radiation in the wild. Science 339: 208–211.

Martin CH & Wainwright PC. 2013c. On the measurement of ecological novelty: scale-eating pupfish are separated by 168 my from other scale-eating fishes. PloS one 8: e71164.

McGirr JA & Martin CH. 2018. Parallel evolution of gene expression between trophic specialists despite divergent genotypes and morphologies. Evolution letters 2: 62–75.

McGirr JA & Martin CH. 2020. Ecological divergence in sympatry causes gene misexpression in hybrids. Molecular ecology 29: 2707–2721.

McGirr JA & Martin CH. 2021. Few Fixed Variants between Trophic Specialist Pupfish Species Reveal Candidate Cis-Regulatory Alleles Underlying Rapid Craniofacial Divergence. Molecular biology and evolution 38: 405–423.

McGlothlin JW, Kobiela ME, Wright HV, Mahler DL, Kolbe JJ, Losos JB & Brodie ED 3rd. 2018. Adaptive radiation along a deeply conserved genetic line of least resistance in Anolis lizards: ADAPTATION AND CONSTRAINT INANOLISLIZARDS. Evolution letters 2: 310–322.

Oksanen J, Blanchet FG, Kindt R, Legendre P, Minchin PR, O’hara RB & Oksanen M. 2013. Package ‘vegan.’ Community ecology package, version 2: 1–295.

Palominos MF, Muhl V & Martin CH. 2024. Tissue-specific transcriptomics uncovers novel craniofacial genes underlying jaw divergence in specialist pupfishes. bioRxiv.

Plath M & Strecker U. 2008. Behavioral diversification in a young species flock of pupfish (Cyprionodon spp.): shoaling and aggressive behavior. Behavioral ecology and sociobiology 62: 1727–1737.

R Core Team R & Others. 2013.R: A language and environment for statistical computing.

Richards EJ, McGirr JA, Wang JR, St John ME, Poelstra JW, Solano MJ, O’Connell DC, Turner BJ & Martin CH. 2021. A vertebrate adaptive radiation is assembled from an ancient and disjunct spatiotemporal landscape. Proceedings of the National Academy of Sciences of the United States of America 118.

Richards EJ & Martin CH. 2017. Adaptive introgression from distant Caribbean islands contributed to the diversification of a microendemic adaptive radiation of trophic specialist pupfishes. PLoS genetics 13: e1006919.

Richards EJ & Martin CH. 2022. We get by with a little help from our friends: shared adaptive variation provides a bridge to novel ecological specialists during adaptive radiation. Proceedings. Biological sciences / The Royal Society 289: 20220613.

Ricklefs RE. 2010. Evolutionary diversification, coevolution between populations and their antagonists, and the filling of niche space. Proceedings of the National Academy of Sciences of the United States of America 107: 1265–1272.

Ripley B, Venables B, Bates DM, Hornik K, Gebhardt A, Firth D & Ripley MB. 2013. Package ‘mass.’ Cran r 538: 113–120.

Roff D. 2000. The evolution of the G matrix: selection or drift? Heredity 84 (Pt 2): 135–142.

Schluter D. 1996. Adaptive radiation along genetic lines of least resistance. Evolution 50.

Schluter D. 2000. The ecology of adaptive radiation. London, England: Oxford University Press.

Schluter D, Clifford EA, Nemethy M & McKinnon JS. 2004. Parallel evolution and inheritance of quantitative traits. The American naturalist 163: 809–822.

Schluter D. 2024. Variable success in linking micro and macroevolution. Evolutionary Journal of the Linnean Society 3: kzae016.

Schmitter-Soto JJ. 1999. Distribution of continental fishes in northern Quintana Roo, Mexico. Southwestern Naturalist 44: 166–172.

Schmitter-Soto JJ. 2017. A revision ofAstyanax(Characiformes: Characidae) in Central and North America, with the description of nine new species. Journal of natural history 51: 1331– 1424.

Schmitter-Soto JJ & Caro CI. 1997. Distribution of tilapia,Oreochromis mossambicus (Perciformes: Cichlidae), and water body characíeristics in Quintana Roo, Mexico. Revista de biologia tropical.

Silva JC e., Potts B & Harrison P. 2020. Population divergence along a genetic line of least resistance in the tree species Eucalyptus globulus. Genes 11.

Simpson GG. 1944. The Major Features of Evolution. Columbia University Press.

St John ME, Dunker JC, Richards EJ, Romero S & Martin CH. 2024. Parallel evolution of integrated craniofacial traits in trophic specialist pupfishes. Ecology and evolution 14: e11640.

Steppan SJ, Phillips PC & Houle D. 2002. Comparative quantitative genetics: evolution of the G matrix. Trends in ecology & evolution 17: 320–327.

Stevenson MM. 1992. Food Habits within the Laguna Chichancanab Cyprinodon (Pisces: Cyprinodontidae) Species Flock. Southwestern Naturalist 37: 337.

Strecker U. 2002. Cyprinodon esconditus, a new pupfish from Laguna Chichancanab, Yucatan, Mexico (Cyprinodontidae). Cybium 26: 301–307.

Strecker U. 2005. Description of a new species from Laguna Chichancanab, Yucatan, Mexico: Cyprinodon suavium (Pisces: Cyprinodontidae). Hydrobiologia 541: 107–115.

Strecker U. 2006a. Genetic differentiation and reproductive isolation in a Cyprinodon fish species flock from Laguna Chichancanab, Mexico. Molecular phylogenetics and evolution 39: 865–872.

Strecker U. 2006b. The impact of invasive fish on an endemic *Cyprinodon* species flock (Teleostei) from Laguna Chichancanab, Yucatan, Mexico. Ecology of freshwater fish 15: 408– 418.

Stroud J, Day JJ, del Rosario Castañeda M & Martin CH. 2025. A global perspective on adaptive radiation: Advances, issues, and future directions. Evolutionary Journal of the Linnean Society.

Stroud JT & Losos JB. 2020. Bridging the Process-Pattern Divide to Understand the Origins and Early Stages of Adaptive Radiation: A Review of Approaches With Insights From Studies of Anolis …. The Journal of heredity.

Svanbäck R & Persson L. 2004. Individual diet specialization, niche width and population dynamics: implications for trophic polymorphisms. Journal of Animal Ecology 73: 973–982.

Tanentzap AJ, Brandt AJ, Smissen RD, Heenan PB, Fukami T & Lee WG. 2015. When do plant radiations influence community assembly? The importance of historical contingency in the race for niche space. The new phytologist 207: 468–479.

Turelli M. 1988. PHENOTYPIC EVOLUTION, CONSTANT COVARIANCES, AND THE MAINTENANCE OF ADDITIVE VARIANCE. Evolution 42.

Wagner C, Harmon L & Seehausen O. 2012. Ecological opportunity and sexual selection together predict adaptive radiation. Nature 487: 366–369.

Wagner CE, Harmon LJ & Seehausen O. 2014. Cichlid species-area relationships are shaped by adaptive radiations that scale with area. Ecology letters 17: 583–592.

Yoder JB, Clancey E, Des Roches S, Eastman JM, Gentry L, Godsoe W, Hagey TJ, Jochimsen D, Oswald BP, Robertson J, Sarver BAJ, Schenk JJ, Spear SF & Harmon LJ. 2010. Ecological opportunity and the origin of adaptive radiations: Ecological opportunity and origin of adaptive radiations. Journal of evolutionary biology 23: 1581–1596.

